# DLDTI: A learning-based framework for identification of drug-target interaction using neural networks and network representation

**DOI:** 10.1101/2020.07.31.230763

**Authors:** Yihan Zhao, Kai Zheng, Baoyi Guan, Mengmeng Guo, Lei Song, Jie Gao, Hua Qu, Yuhui Wang, Ying Zhang, Dazhuo Shi

## Abstract

To elucidate novel molecular mechanisms of known drugs, efficient and feasible computational methods for predicting potential drug-target interactions (DTI) would be of great importance. A novel calculation model called DLDTI was generated for predicting DTI based on network representation learning and convolutional neural networks. The proposed approach simultaneously fuses the topology of complex networks and diverse information from heterogeneous data sources and copes with the noisy, incomplete, and high-dimensional nature of large-scale biological data by learning low-dimensional and rich depth features of drugs and proteins. Low-dimensional feature vectors were used to train DLDTI to obtain optimal mapping space and infer new DTIs by ranking DTI candidates based on their proximity to optimal mapping space. DLDTI achieves promising performance under 5-fold cross-validation with AUC values of 0.9172, which was higher than that of the method based on different classifiers or different feature combination technique. Moreover, biomedical experiments were also completed to validate DLDTI’s performance. Consistent with the predicted result, tetramethylpyrazine, a member of pyrazines, reduced atherosclerosis progression and inhibited signal transduction in platelets, via PI3K/Akt, cAMP and calcium signaling pathways. The source code and datasets explored in this work are available at https://github.com/CUMTzackGit/DLDTI

## Introduction

Research on drug development is becoming increasingly expensive, while the number of newly approved drugs per year remains quite low [1][2]. In contrast to the classical hypothesis of “one gene, one drug, one disease”, drug repositioning aims to identify new characteristics of existing drugs [3]. Considering the available data on safety of already-licensed drugs, this approach could be advantageous compared with traditional drug discovery, which involves extensive preclinical and clinical studies [4]. Currently, a number of existing drugs have been successfully tuned to the new requirements. Methotrexate, an original cancer therapy, has been used for the treatment of rheumatoid arthritis and psoriasis for decades [5]. Galanthamine, an acetylcholinesterase inhibitor for treating paralysis, has been approved for Alzheimer’s disease [6].

Besides the evidence based on biological experiments and clinical trials, computational methods could facilitate high-throughput identification of novel target proteins of known drugs. To discover targets of drugs with known chemical structures, the prediction of drug-target interaction (DTI) based on numerous computational approaches have provided an alternative to costly and time-consuming experimental approaches [7]. In the past years, DTI prediction has bolstered the identification of putative new targets of existing drugs [8]. For instance, the computational pipeline predicted that telmisartan, an angiotensin II receptor antagonist, had the potential of inhibiting cyclooxygenase. In vitro experimental evidence also validated the predicted targets of this known drug [9]. Further, combined with in silico prediction, in vitro validation and animal phenotype model demonstrated that, topotecan, a topoisomerase inhibitor also had the potential to act as a direct inhibitor of human retinoic-acid-receptor-related orphan receptor-gamma t (ROR-γt) [10].

Most existing prediction methods mainly extract information from complex networks. Bleakley et al. [11] proposed a support vector machine-based method for identifying drug-target interactions based on bipartite local model (BLM). Mei et al. [12] proposed BLMNII method for predicting DTIs based on the bipartite local model and neighbor-based interaction-profile inference. In addition, some researchers adopted kernelized Bayesian matrix factorization to predict DTIs, called KBMF2K [13]. A key step of KBMF2K is utilizing dimensional reduction, matrix factorization, and binary classification. Although homogenous network-based derivation methods have achieved good results, they are less effective in low-connectivity (degree) drugs for known target networks. The introduction of heterogeneous information can provide more perspective for predicting the potential of DTI. Recently, Luo et al. proposed a heterogeneous network-based unsupervised method for computing the interaction score between drugs and targets, called DTInet [9]. Subsequently, they proposed a neural network-based method [14] for improving the prediction performance of DTI. Effective integration of large-scale heterogeneous data sources is crucial in academia and industry.

Tetramethylpyrazine (TMPZ) is a member of pyrazines derived from Rhizoma Chuanxiong Hort [15]. According to a recent review, TMPZ could attenuate atherosclerosis by suppressing lipid accumulation in macrophages [16], alleviation of lipid metabolism disorder [17], and attenuation of oxidative stress [18]. However, since atherosclerosis is a chronic illness involving multiple cells and cytokines [19], besides lipoprotein metabolism and oxidative stress, other possible targets of TMPZ on atherosclerosis remain unexplored.

In this study, a novel model for prediction of DTI based on network representation learning and convolutional neural networks, referred to as DLDTI is presented for in silico identification of target proteins of known drugs. New DTIs were inferred by integrating drug- and protein-related multiple networks, to demonstrate the DLDTI’s ability of integrating heterogeneous information and neural networks to extract deep features of drugs and target networks as well as attributes to effectively improve prediction accuracy. Moreover, comprehensive testing demonstrated that DLDTI could achieve substantial improvements in performance over other prediction methods. Based on the results predicted by DTDTI, new interactions between TMPZ and targets involved in atherosclerosis, namely signal transduction in platelets, were validated in vivo. The anti-atherosclerosis effect of TMPZ was confirmed in a novel atherosclerosis model. In summary, these improvements could advance studies on drug-target interaction.

## Results

### Overview of DLDTI and performance evaluation on predicting drug-target interaction

A new computational model referred to as DLDTI was developed to predict potential DTIs to identify novel behavior of traditional drugs based on complex networks and heterogeneous information. As an overview (Figure 1), DLDTI integrates learning from complex network’s various heterogeneous information to obtain low-dimensional and deep rich features (Figure 2), through a processing method known as compact feature learning. During compact feature learning, the resulting low-dimensional descriptor integrates attribute characteristics, interaction information, relational properties, and network topology of each protein or target node in the complex network. DLDTI then determines the optimal mapping from the plenary mapping space to the prediction subspace, and whether the feature vector is close to the known correlations. Afterwards, DLDTI infers the new DTIs by ranking the drug-target interaction candidates according to their proximity to the predicted subspace.

**Figure 1.**
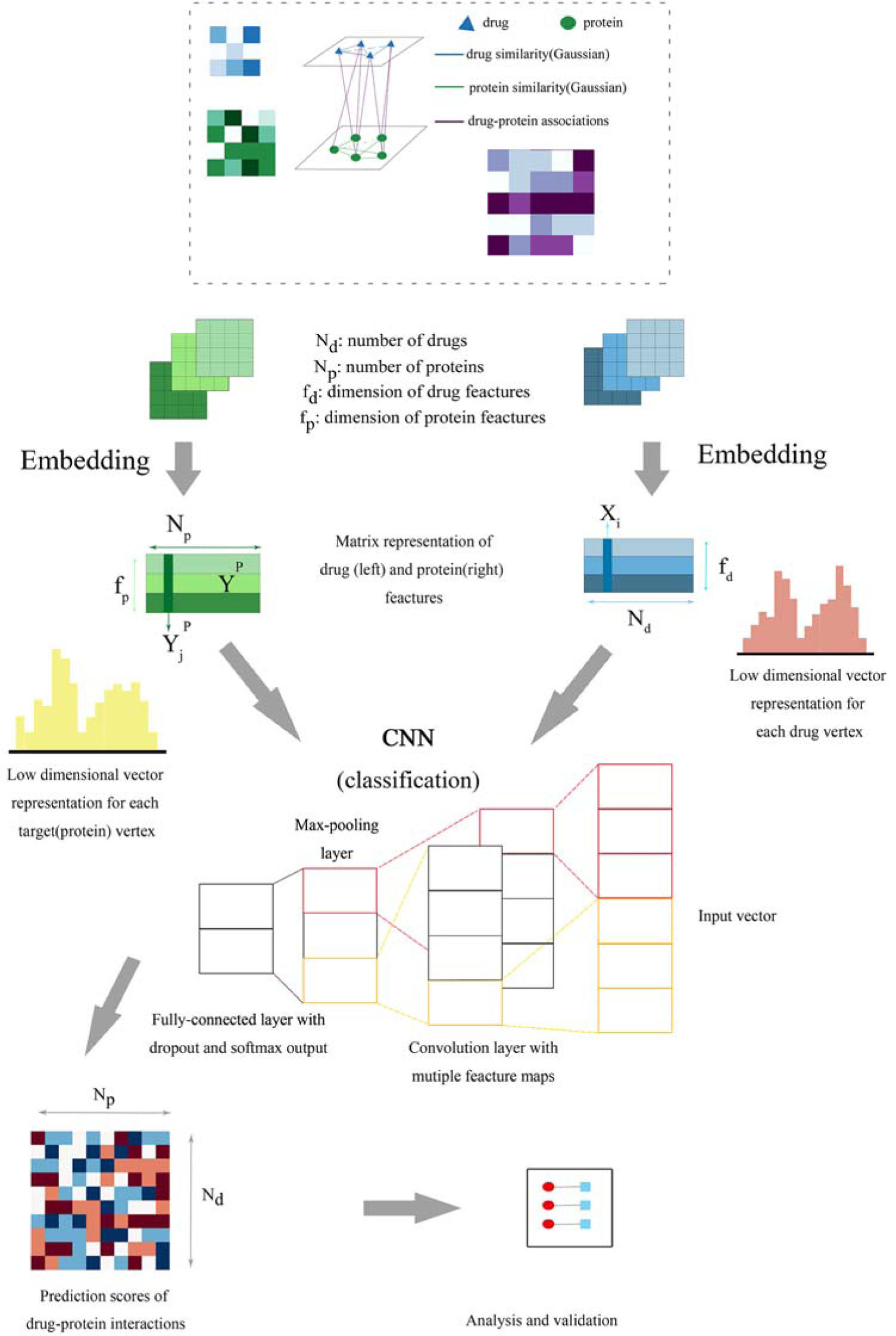
The flowchart of the DLDTI pipeline. DLDTI first integrates a variety of drug-related information sources to construct a heterogeneous network and applies a compact feature learning algorithm to obtain a low-dimensional vector representation of the features describing the topological properties for each node. Next, DLDTI determines the optimal mapping from the plenary mapping space to the prediction subspace, and whether the feature vector is close to the known correlations. Afterwards, DLDTI infers the new DTIs by ranking the drug-target interaction candidates according to their proximity to the predicted subspace

**Figure 2.**
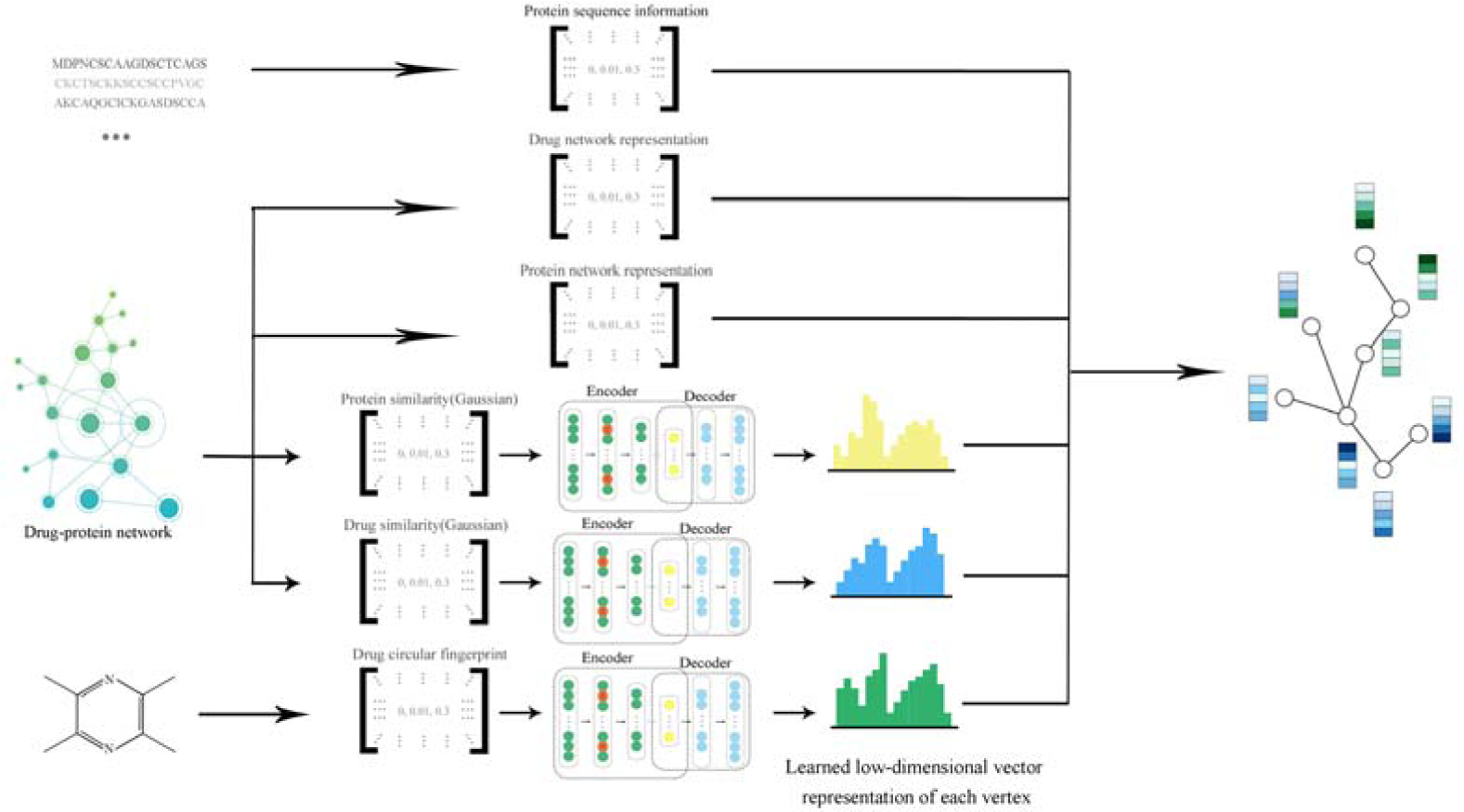
Schematic illustration of compact feature learning. The Node2Vec algorithm is firstly used to calculate the topology information in complex networks. Gaussian interaction profile kernel similarity (GIP) and drug structure information are then extracted by a stacked automatic encoder, and the heterogeneous information is integrated to obtain a low-dimensional representation of the feature vector of each node. The resulting low-dimensional descriptor integrates the attribute characteristics, interaction information, relationship attributes and network topology of each protein or target node in the complex network.

DLDTI yields accurate DTI prediction. Firstly, the predictive performance of DLDTI was assessed using five-fold cross-validation, where randomly selected subset of one-fifth of the validated drug-target interaction were paired with an equal number of randomly sampled non-interacting pairs to derive the test set. The remaining 75% of known drug-target interaction and same number of randomly sampled non-interacting pairs were used to train the model. DLDTI was compared with three methods based on different classifiers used for DTI prediction, including DTI-ADA, DTI-KNN, and DTI-RF [20][21][22]. The comparison revealed that DLDTI consistently outperforms the other three methods, with 0.93% higher AUC, 3.55% higher AUPR, 0.61% higher accuracy (Acc), 3.96% higher precision (Pre) than the second-best method (Fig. 3c, Fig. 3d and Fig. 3e). Compared to DTI-ADA (which predicts DTI based on the AdaBoost classifier), the DLDTI of the AUROC and AUPR was 6.96% and 7.81% higher, respectively, which could have been due to the inability of traditional machine learning to extract deeper abstract features for prediction, resulting in poor performance, while DLDTI applies a deep convolutional neural network approach and is able to capture the potential structural properties of complex networks and heterogeneous information.

**Figure 3.**
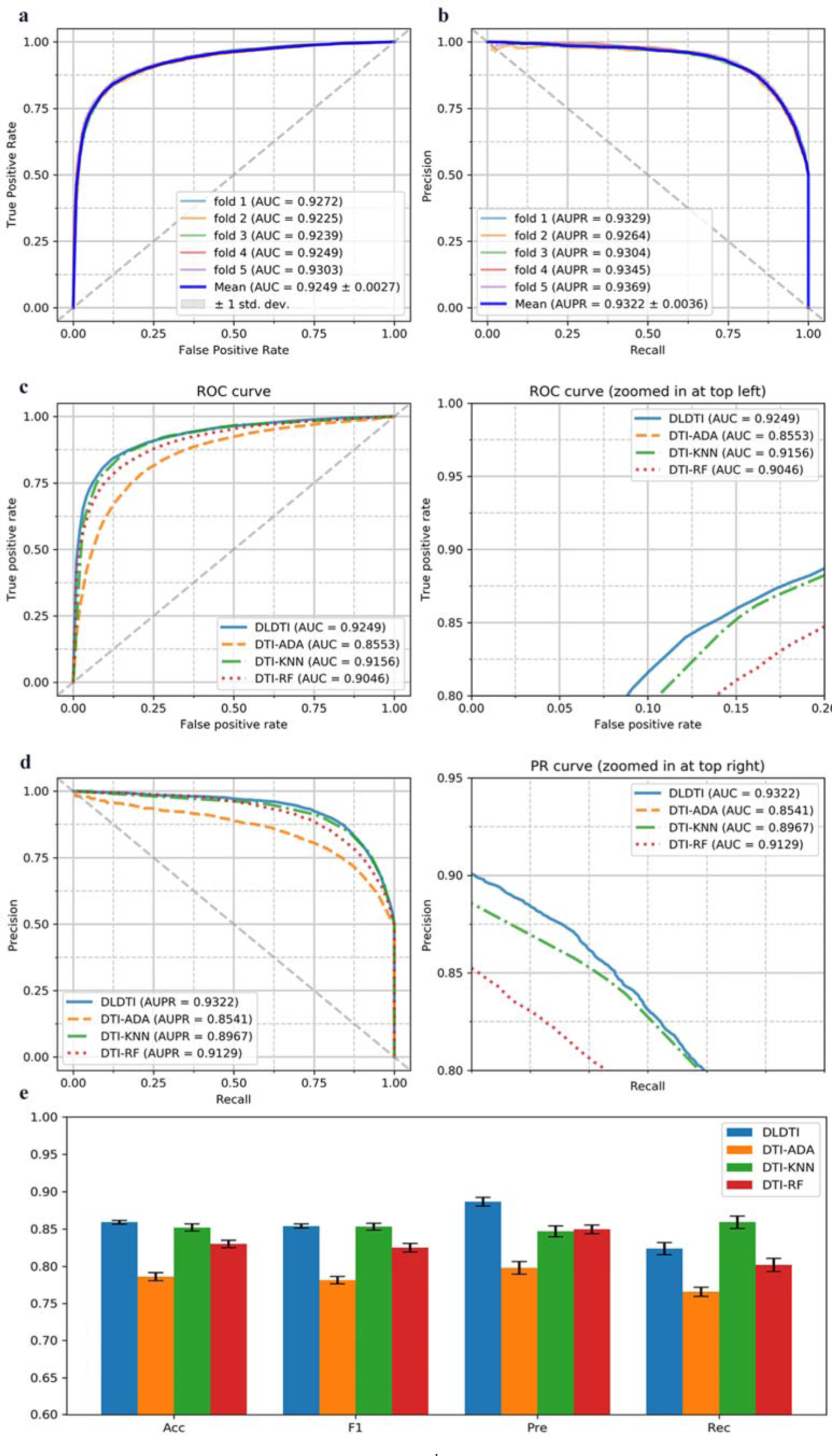
Performance of DLDTI. (a) ROC curves performed by DLDTI model on DrugBank dataset. (b) PR curves performed by DLDTI model on DrugBank dataset. (c) Performance comparison (AUC scores) among four different prediction model which are DTI-ADA, DTI-KNN, and DTI-RF.(d)Performance comparison (AUPR scores) among four different prediction models including DTI-ADA, DTI-KNN, and DTI-RF.(e)Performance comparison (Acc., F1, Pre., Rec. scores) among DTI-ADA, DTI-KNN, and DTI-RF prediction models.

### Enrichment analysis suggested that TMPZ might affect signal transduction pathways involved in platelet activation

To elucidate the potential function of TMPZ on atherosclerosis, the predicted results from DLDTI model were uploaded to the search tool for retrieval of interacting genes/proteins database (STRING, Version 11) (https://string-db.org/) [23] to determine over-represented Kyoto Encyclopedia of Genes (KEGG) pathways and Genomes Gene Ontology (GO) categories. GO analysis demonstrated that 31.4% of genes were involved in signal transduction (Supplemental Table 1). As shown in Table 1, phosphoinositide 3-kinase (PI3K)/Akt signaling pathway, neuroactive ligand-receptor interaction, mitogen-activated protein kinase (MAPK) signaling pathway, calcium signaling pathway, repressor activator protein 1 (Rap1) signaling pathway, cyclic guanosine monophosphate (cGMP)-protein kinase G (PKG) signaling pathway, and cyclic adenosine monophosphate (cAMP) signaling pathway were the top-ranked results of KEGG enrichment. It is noteworthy that ADP-mediated platelet activation via purinergic receptors included almost all signal transduction pathways shown in Table 1 [24][25]. Interestingly, among the 288 predicted targets of TMPZ on atherosclerosis, 190 proteins were also involved in the platelet activation process (Supplemental Table 2). Therefore, it was assumed that the anti-atherosclerosis potential of TMPZ could be largely attributed to its inhibition of purinergic receptor-dependent platelet activation, which involves signal transduction pathways such as PI3K/Akt. Based on the predicted result, clopidogrel, an anti-platelet drug widely used in the clinical application, was chosen as the positive control.

**Table 1.**
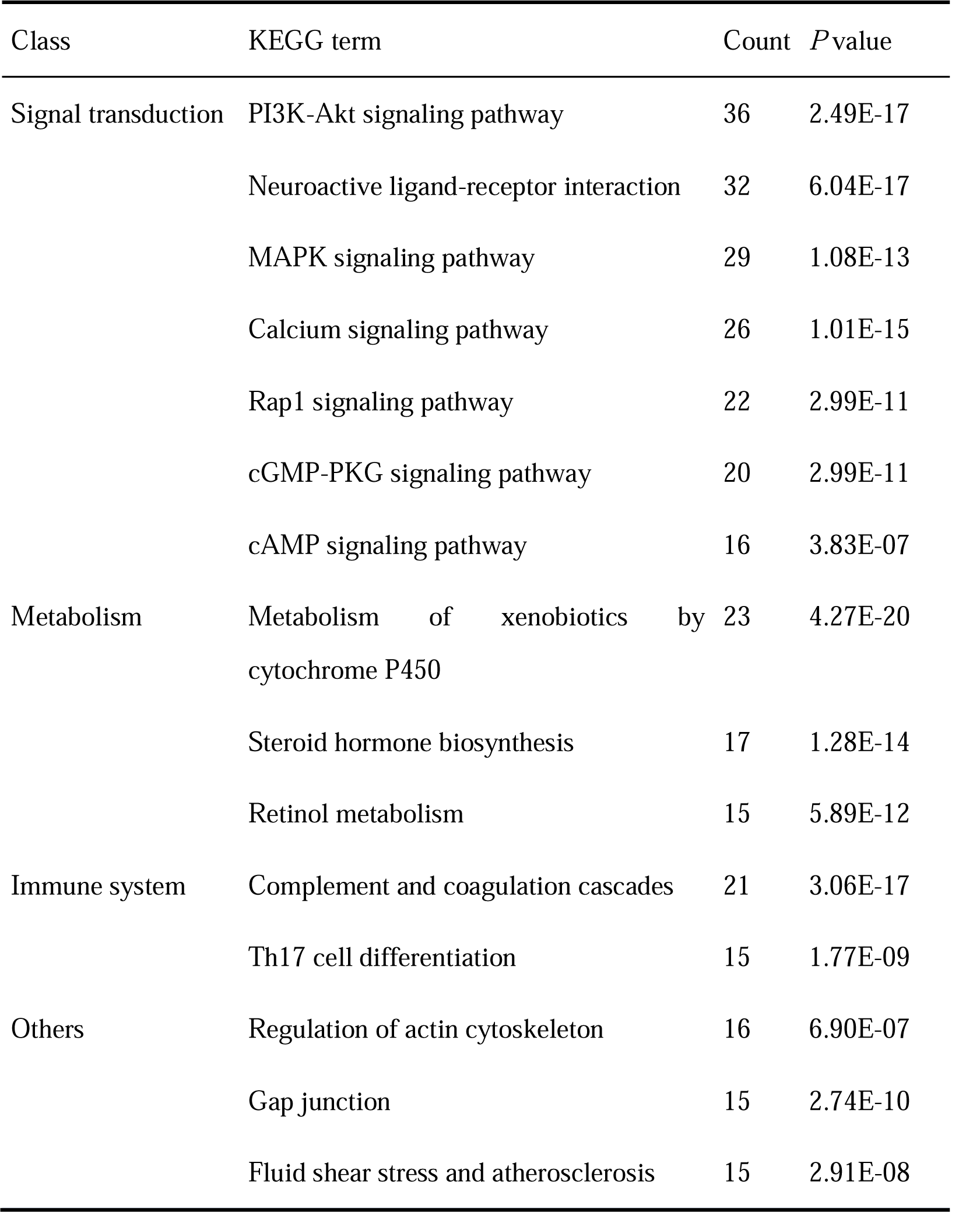
KEGG pathway enrichment analysis of DLDTI results

| Class | KEGG term | Count | *P* value |
| --- | --- | --- | --- |
| Signal transduction | PI3K-Akt signaling pathway | 36 | 2.49E-17 |
|  | Neuroactive ligand-receptor interaction | 32 | 6.04E-17 |
|  | MAPK signaling pathway | 29 | 1.08E-13 |
|  | Calcium signaling pathway | 26 | 1.01E-15 |
|  | Rap1 signaling pathway | 22 | 2.99E-11 |
|  | cGMP-PKG signaling pathway | 20 | 2.99E-11 |
|  | cAMP signaling pathway | 16 | 3.83E-07 |
| Metabolism | Metabolism of xenobiotics by cytochrome P450 | 23 | 4.27E-20 |
|  | Steroid hormone biosynthesis | 17 | 1.28E-14 |
|  | Retinol metabolism | 15 | 5.89E-12 |
| Immune system | Complement and coagulation cascades | 21 | 3.06E-17 |
|  | Th17 cell differentiation | 15 | 1.77E-09 |
| Others | Regulation of actin cytoskeleton | 16 | 6.90E-07 |
|  | Gap junction | 15 | 2.74E-10 |
|  | Fluid shear stress and atherosclerosis | 15 | 2.91E-08 |

### Validation

#### Ldlr-/- hamsters developed severe hyperlipidemia and atherosclerosis lesions when fed with HFHC diet

Before dietary induction, genotypes were determined by PCR analysis. Using ear genomic DNA, 194-nucleotide deletion (Δ194) was detected in homozygous (-/-) hamsters (Figure 4a). After feeding them on high-fat and high-cholesterol (HFHF) diet for 16 weeks, low-density lipoprotein receptor knock-out (Ldlr-/-) hamsters developed severe hyperlipidemia. As an antiplatelet medication, clopidogrel did not influence circulating levels of Total cholesterol (TC), triglyceride (TG), high-density lipoprotein (HDL) and non-HDL (Figure 4b, 4c, 4d and 4e). Compared with vehicle-treated hamsters, decreased levels of TC (*p*<0.05) and non-HDL (*p*<0.05) were observed in TMPZ-treated group (Fig. 4b and 4d). However, TMPZ did not influence TG or HDL levels.

**Figure 4.**
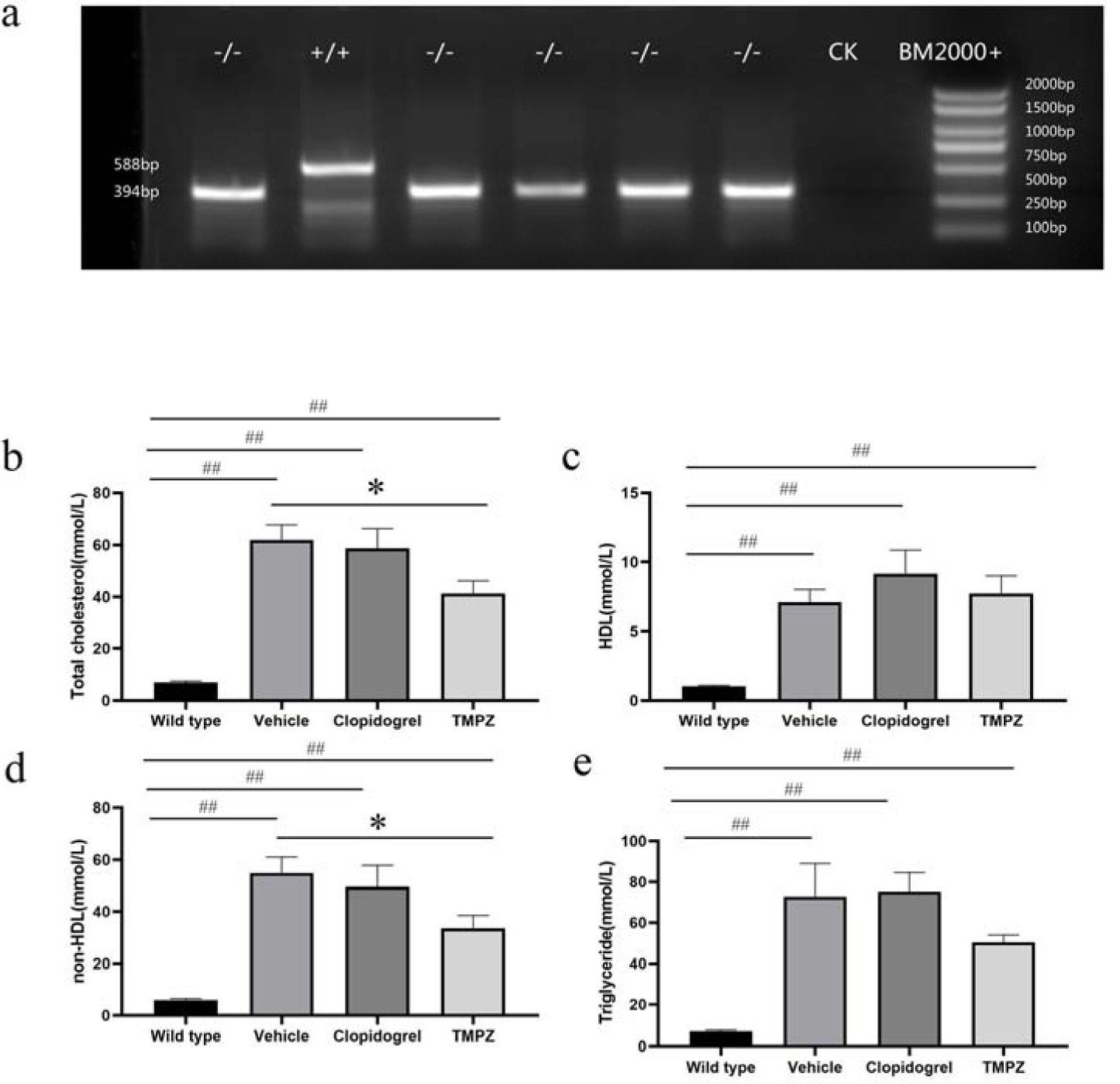
Genotyping and lipid parameters between different groups. (a).PCR analysis was performed using ear genomic DNA from WT (+/+) and homozygote (−/−) with the Δ194 deletion. The concentrations of plasma TC (b), HDL(c), non-HDL(d) and TG(e) were measured in WT, vehicle, TMPZ and clodipogrel groups at the endpoint of this experiment. Differences were assessed by unpaired student t’s test or Mann-Whitney test. * *p*<0.05 versus Vehicle, \*\**p*<0.01 versus Vehicle. ##*p*<0.01 versus WT. All data was expressed as mean ±standard error (SEM)

#### TMPZ ameliorated atherosclerosis lesion progression

The *en face* analysis demonstrated that vehicle-treated hamsters developed significant atherosclerotic lesions (mean value 28.38%) throughout the whole aorta. However, atherosclerotic lesions induced by the same dietary manipulation in TMPZ- and clopidogrel-treated groups were significantly decreased (mean value 10.02% and mean value 17.47%, respectively) (Figure 5a and 5b). It’s noteworthy that the lesion area in TMPZ-treated group was also less than that in clopidogrel-treated group (Figure 5b). As the blank control group, WT hamsters on chow diet did not develop any lesions throughout the aorta.

**Figure 5.**
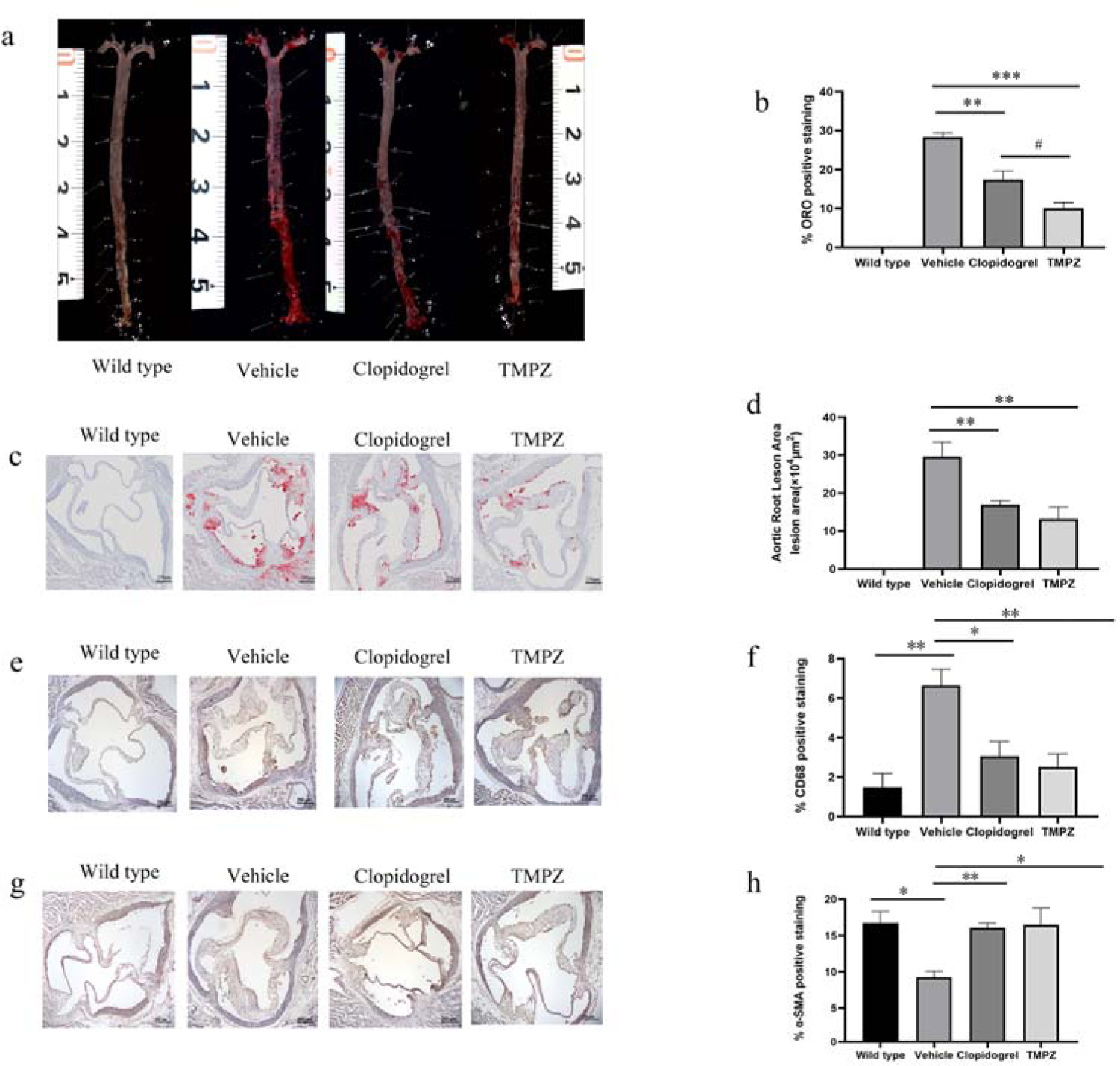
Histological analysis. (a) Representative images of en face analysis. (b) Quantitative analysis of lesion areas in whole aortas. Differences were assessed by unpaired student t’s test. (c) Representative images of ORO staining of aortic root sections. (d) Quantitative analysis of lesion areas in aortic root sections. (e) Representative images of macrophage (CD68) analysis (b) Quantitative analysis of lesions area in macrophage analysis. (f) Representative images of SMC (SMA) analysis (g) Quantitative analysis of lesions area in SMC. Differences were assessed by unpaired student t’s test. * *p*<0.05 versus Vehicle, \*\**p*<0.01 versus Vehicle. #*p*<0.05 versus clopidogrel. Scale bar=250μm. All data was expressed as mean ± SEM.

Similar to the *en face* analysis, the HFHC fed vehicle group had significantly increased lesion areas (mean area 29.58×10^4^ μm^2^) in aortic roots compared to the blank controls measured by image analysis of Oil Red O staining, and either TMPZ (mean area 13.25×10^4^ μm^2^) or clopidogrel (mean area 16.99×10^4^ μm^2^) treatment reduced the lipid-rich areas (Figure 5c and 5d).

Under the stimulation of adhesion molecules, monocytes infiltrate into the intima and differentiate into macrophages [26]. Besides macrophage accumulation, diminished smooth muscle cells (SMC) could also exacerbate the formation of unstable plaques [27]. To determine the components of atherosclerosis lesions in the aortic root, immunohistochemical staining for macrophages and SMC was performed [28]. Histopathological evaluation of macrophages accumulation revealed differences in CD68-positive areas between the groups. As shown in Figure 5e and 5f, the percentage of macrophage positive staining in lesions was increased by atherosclerosis progression in the vehicle-treated group. WT group (mean value 1.48%) had significantly fewer macrophage accumulation than vehicle-treated group (mean value 6.65%). Infiltrated macrophages in lesions were significantly decreased by TMPZ (mean value 2.52%) or clopidogrel (mean value 3.07%) treatment. As shown in Figure 5g and 5h, besides macrophage infiltration, the percentage of a-SMA positive staining was diminished in Ldlr-/- hamsters (mean value 9.27%) compared with the WT hamsters (mean value 16.76%). Administration TMPZ (mean value 16.50%) or clopidogrel (mean value 16.09%) for 8 weeks could ameliorate SMC reduction in atherosclerosis lesions.

#### TMPZ inhibited signaling transduction in ADP-mediated platelet activation

In addition to the surrogates of platelet activation, calcium and cAMP signaling are also essential in signal transduction. Downstream from Gq signaling, protein kinase C (PKC) activation results in the formation of inositol triphosphate (IP3), which leads to an elevation of intracellular calcium [24]. Calcium mobilization is also required for the phosphorylation of Akt (also known as protein kinase B) in PI3K/Akt signaling pathway [29]. In response to ADP, Gi signaling activation mediates the inhibition of AC, resulting in the diminished synthesis of cAMP. The inhibitory effect of Gi on cAMP synthesis could cause platelet activation [25].

Figure 6 shows that fura-2/AM is a membrane-permeant calcium indicator. The ratio of F340/F380 is directly correlated to the amount of intracellular calcium. The data revealed that TMPZ and clopidogrel markedly inhibited calcium mobilization, as detected using fluorescence mode of Synergy H1 microplate reader. Moreover, TMPZ-and clopidogrel-treated groups showed a higher concentration of cAMP in the active platelets. These findings indicate that TMPZ and clopidogrel could inhibit calcium mobilization and elevate intracellular concentration of cAMP, thereby inhibiting platelet activation.

**Figure 6.**
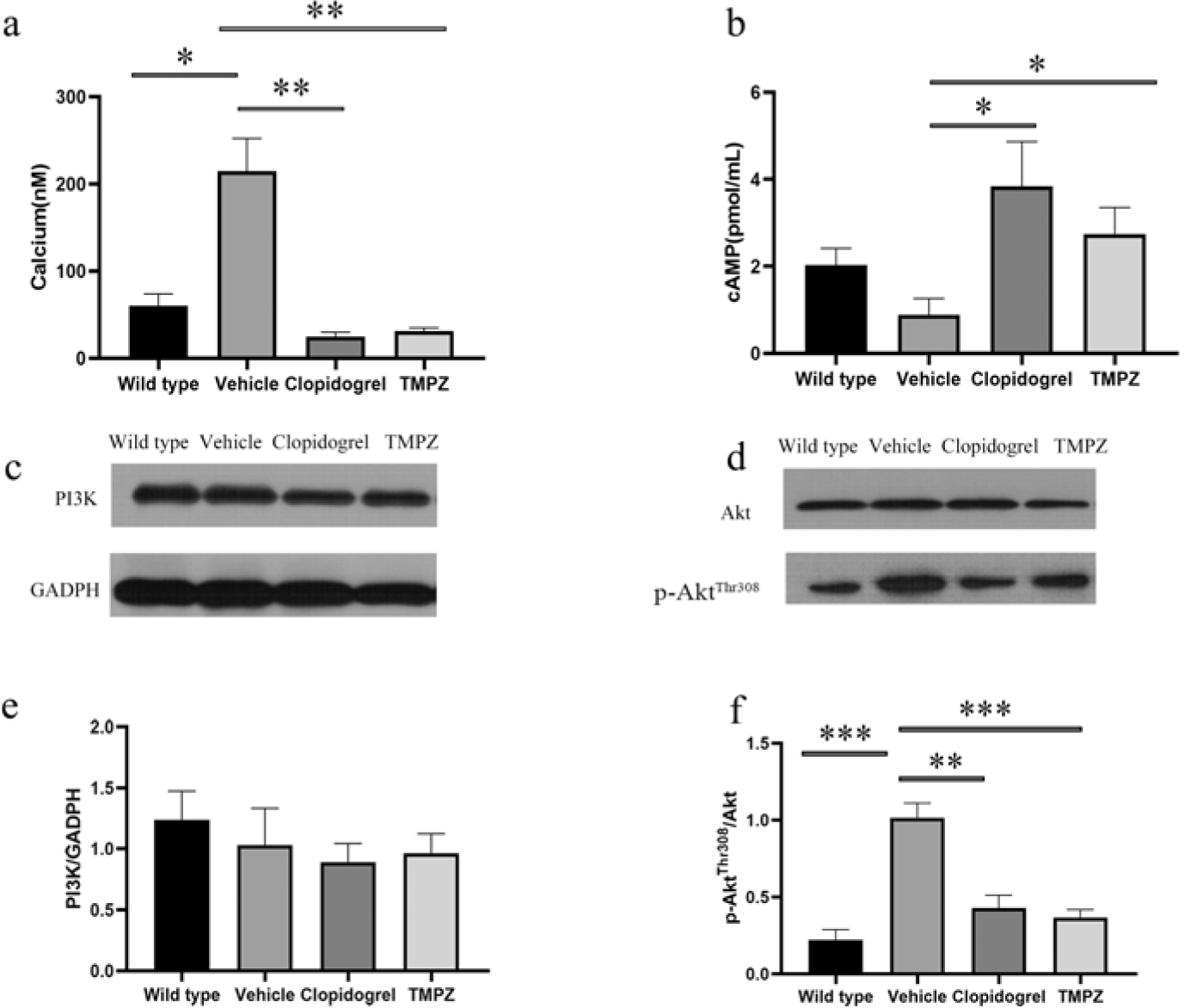
Signaling transduction in ADP-mediated platelet activation. (a) Intracellular calcium concentration. (b) Intracellular cAMP concentration. Western blot analyses of the expression of PI3K (c), Akt (d) and p-Akt (d). Differences were assessed by unpaired student t’s test with or without Welch’s corrections. ** *p*<0.01 versus Vehicle, * *p*<0.05 versus Vehicle. All data was expressed as mean ±SEM.

As the major downstream effector of PI3K, Akt plays an essential role in the regulation of platelet activation. Stimulation of platelets with ADP could result in Akt activation, which was indicated by Akt phosphorylation [29]. The protein expressions of PI3K, Akt, and p-Akt in the top-ranked signal transduction pathway were measured to validate the predicted pathways. ADP-induced P2Y12 receptor activation could cause PI3K dependent Akt phosphorylation, a critical positive regulator pathway for signal amplification. There was no difference in PI3K expression levels between WT, vehicle, TMPZ, and clopidogrel groups (Figure 6c). Phosphorylation of Akt was inhibited by TMPZ or clopidogrel administration when compared with vehicle-treated group. It is noteworthy that phosphorylation of Akt did not differ between WT, TMPZ and clopidogrel groups, which indicates that platelet activity in atherosclerosis hamsters treated with TMPZ or clopidogrel could be comparable to that in healthy ones (Figure 6d). These findings indicate that TMPZ and clopidogrel could attenuate Akt signaling, thereby blocking the platelet activation induced by ADP.

## Discussion

In summary, this study provides a novel DTI model and validates its efficacy via a novel atherosclerosis model. This DLDTI model could provide an alternative to the high-throughput screening of drug targets. The proposed approach simultaneously fuses the topology of complex networks and diverse information from heterogeneous data sources and copes with the noisy, incomplete, and high-dimensional nature of large-scale biological data by learning the low-dimensional and rich depth features of drugs and proteins. The low-dimensional descriptors learned by DLDTI capture attribute characteristics, interaction information, relational properties, and network topology attributes of each drug or target node in a complex network. The low-dimensional feature vectors were used to train DLDTI to obtain the optimal mapping space and to infer new DTIs by ranking drug-target interaction candidates based on their proximity to the optimal mapping space. New DTIs were inferred by integrating drug- and protein-related multiple networks, to demonstrate DLDTI’s ability to integrate heterogeneous information and that deep neural networks are capable of extracting drug and target networks and that deep features of attributes can effectively improve the prediction accuracy. This work also proved that TMPZ administration could attenuate atherosclerosis lesions, characterized by diminished lipid deposition, macrophage accumulation, and increased SMC percentage. Moreover, TMPZ could inhibit platelet activation by inhibiting Akt’s phosphorylation and calcium mobilization and increasing intracellular cAMP concentration.

The current study proposes a learning-based framework called DLDTI for identifying the association of drug targets. The structural characteristics of drug and the characteristics of the protein properties were firstly extracted. An automatic encoder-based model was then proposed for feature selection. Using this feature representation, a convolutional neural network architecture was proposed for predicting the DTI. The advantages of DLDTI were demonstrated by comparing it with three different methods. Experiments on DTI showed that the performance of DLDTI was better than that of the alternative method, which shows that the proposed learning-based framework was properly designed.

Furthermore, in the validation study of the DLDTI model, we used TMPZ (a drug with known structure) to explore its effects on atherosclerosis in vivo. Consistent with previous studies [16][17][18], the results revealed that TMPZ could ameliorate the phenotyping of atherosclerosis in Ldlr-/- hamsters, a novel atherosclerosis model [30][31]. Diminished lipid deposition and macrophage accumulation, and increased percentage of SMC were observed in TMPZ- and clopidogrel-treated hamsters, in comparison with vehicle-treated animals. Interestingly, it was found that the majority of potential pathways of TMPZ on atherosclerosis were also involved in signal transduction of platelet activation. From the initial endothelial dysfunction in the early stage to the destabilized plaques in the advanced stage, platelet plays a pivotal role [32]. Activated platelets act as the key trigger for rupture-prone plaque formation. Current evidence shows that platelet hyperactivity is associated with a prothrombotic state and increases the incidence of recurrent cardiovascular events among patients with coronary artery disease [33]. Over the past decade, it has been found that platelets can be activated by various stimuli like collagen, thrombin, and ADP. Based on the pathway analysis of predicted results, this work focused on signal transduction in ADP-mediated platelet activation (Table 1). The results revealed that the activated signal transductions, characterized by increased calcium mobilization, decreased cAMP concentration and increased phosphorylation of Akt were observed in ex vivo platelets from vehicle-treated hamsters. Platelets from TMPZ- and clopidogrel-treated hamsters showed increased cAMP level and diminished calcium mobilization and phosphorylation of Akt.

Future studies will focus on solving “cold-start” problem, which is faced by all algorithms that apply collaborative filtering technology. In the current study, the top three feature vectors with the highest scores are weighted by 60%, 30%, and 10%, respectively, based on the similarity of protein sequences and the similarity of drug structures, to obtain new interaction feature vectors to solve the cold start problem. In addition, to validate the study, the top-ranked pathways of signal transduction involved in platelet activation were examined, although reduced TC and non-HDL levels and diminished macrophage accumulation in lesions were also observed. These effects also could also contribute to the diminished total lesions area and be the topic of our following research.

## Materials and Methods

### Prediction experiments

#### Human drug-target interactions database

The current study used the DrugBank (http://www.drugbank.ca) established by Wishart *et al*. as the benchmark [34]. The chemical structure of each drug in SMILES format was extracted from the DrugBank. This study used drugs that satisfied the human target represented by a unique EnsemblProt login number. In summary, 904 drugs and 613 unique human targets (proteins) were linked to construct a drug-target interaction network of positive samples, while a matching number of unknown drug-target pairs (by excluding all known DTIs) was randomly selected as negative samples.

### Feature representation

#### Gaussian interaction profile kernel similarity for drugs and targets

According to previous studies, drug similarity can be determined by calculating their nuclear similarity through Gaussian interaction profile kernel similarity (GIP) [35][36]. The GIP similarity between drug *d*_*i*_ and drug *d*_*j*_ is defined as follow:

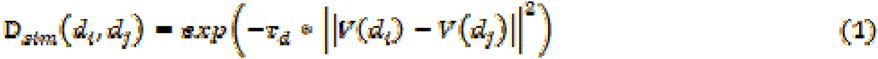

Where, the binary vector *V*(*d*_*i*_) and *V*(*d*_*j*_) are the *i*-th and *j*-th row vectors of the drug-target interaction network *A*. .τ_*d*_ is the kernel bandwidth and is computed by normalizing the original parameter .τ_*d*_*′*:

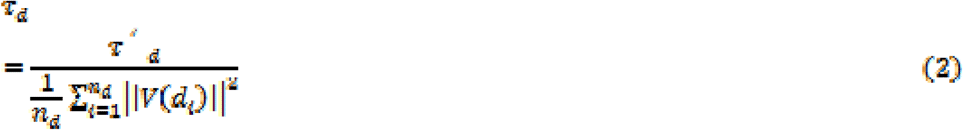

Similarly, the GIP similarity for targets can be defined as follows:

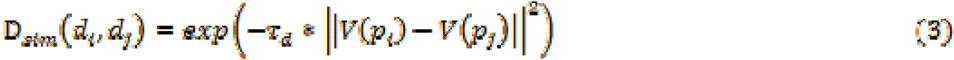

Where, the binary vector *V*(*p*_*i*_) and *V*(*p*_*j*_) are the *i*-th row and the *j*-th column vector of the drug-target interaction network *A*, respectively.τ_*p*_ is the kernel bandwidth and is computed by normalizing the original parameter .τ_*p*_*′*:

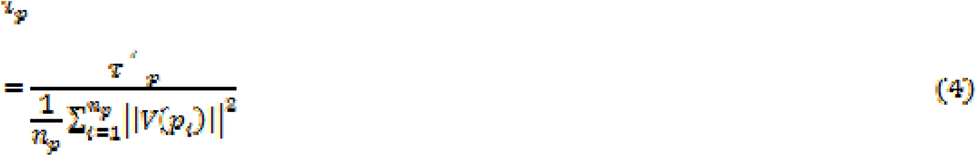

#### Protein sequence feature

The sequences for drug targets (proteins) in Homo sapiens were downloaded from STRING. The *k-mer* algorithm was used to count Subsequence information in protein sequences and used as a feature vector to solve alignment issues presented by differences in sequence length [37].

#### Drug structure feature

Morgan and circular fingerprints were used to map the structure information of drugs to feature vectors based on SMILES for drugs downloaded from the DrugBank database.

#### Graph embedding-based feature for drugs and targets

Graph data is rich in behavioral information about nodes, which can be used as a comprehensive descriptor for drugs and targets [38]. To map a high-dimensional dense matrix like graph data to a low-density vector, a Graph Factorization algorithm [39] was hereby introduced. Graph factorization (GF) is a method for graph embedding with time complexity O(|E|). To obtain the embedding, GF factorizes the adjacency matrix of the graph to minimize loss functions as follow:

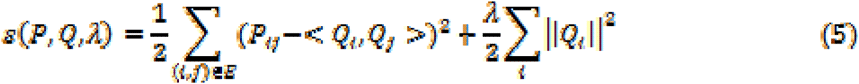

Where, λ is the regularization coefficient.*P* and *Q* are the adjacency matrix with weights and factor matrix, respectively. *E* is the set of edges, which includes *i* and *j*.

The gradient of the function ε with respect to *Q*_*i*_ is defined as follow:

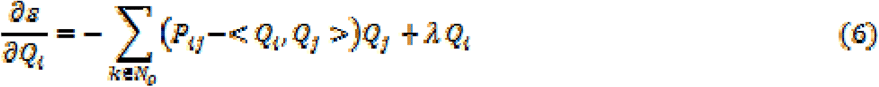

Where, *N*_*o*_ is the set of neighbors of node *o* and the Graph Factorization algorithm, graph embeddings and targets in the drug-target interaction network can be obtained to describe their behavioral information.

#### Stacked Autoencoder

Since DLDTI integrates heterogeneous data from multiple sources, including protein sequence, drug structure, and drug-target interaction network information, the integrated biological data is characterized by noise, incompleteness and has high-dimension. Therefore, stack autoencoder (SAE) was used to establish the optimal mapping of drug space to target space to obtain low dimensional drug Feature vector [40][41]. SAE can be defined as follows:

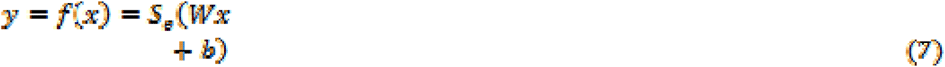

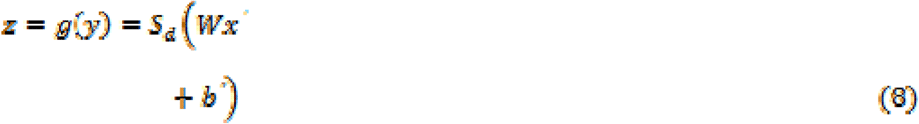

Where *y* and *z* are encoding and decoding function, respectively. *W* and *W′* are the relational parameters between two layers, respectively. *b* and *b′* are vectors of bias parameters. The activation function used is ReLU:

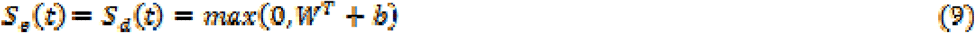

#### Convolutional neural network

Convolutional neural networks were proposed by Lecun *et al*. in 1989[42]. Subsequently, they have performed well in image classification, sentence classification, and biological data analysis. In this study, convolutional neural networks were used to train supervised learning models to predict potential drug-target interactions. They were also chosen as supervised learning models to study deep features and predict potential drug targets interaction. The model used has convolutional and activation, Maxpooling, fully connected and softmax layers. Their roles are to extract depth features, down-sample, and classify samples, respectively. The convolutional layer is one of the most important parts of the CNN and aims to learn the deep characteristics of the input vectors, which is defined as follows;

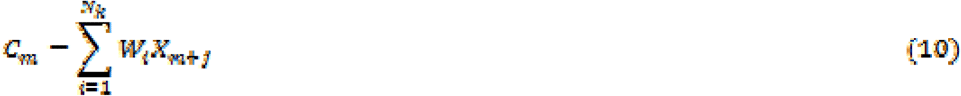

Where; *X* is the input feature of length *L, N*_*k*_ is the number of kernels, *m*={*0,…,L − N*}, and W is a weight vector of length *N*_*k*_. The feature map *C*_*m*_ is then put into the activation function ReLU, which is defined as follow:

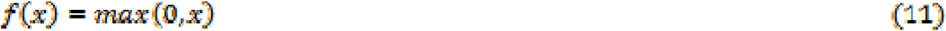

The ReLU function increases the nonlinear relationship between the layers of the neural network, saves computation, solves the problem of gradient disappearance, and reduces the interdependence of parameters to mitigate the problem of overfitting.

The convolutional and maximum pooling layers can extract important features from the input vectors. The output of all kernels was then concatenated into a vector and fed to the fully-connected layer *f*(*w* · *y*). Where; *y* is the output of Maxpooling layer and *w* is the weight matrix. Finally, the softmax layer scored the input vectors as a percentage.

#### Pathway analysis of predicted results from DLDTI

Atherosclerosis-related gene sets were downloaded from GeneCards (https://www.genecards.org/) [43]. After using retrieve tool on Uniprot database (https://www.uniprot.org/), different identifiers from Drug Bank and GeneCards were converted to UniProtKB. Based on intersection of potential targets of TMPZ from DLDTI model and confirmed target proteins of atherosclerosis, the matched targets were regarded as the predicted targets of TMPZ on atherosclerosis. The predicted targets were uploaded to STRING for KEGG pathway and GO analysis.

### Validation experiments

#### Ldlr-/- hamsters

This study was approved by the Animal Ethics Committee of Xiyuan Hospital and strictly adhered to the principles of laboratory animal care (NIH publication No.85Y23, revised 1996). Male, 8-week aged and Ldlr-/- hamsters were provided by the health science center, Peking University. The Ldlr-/- genotype was confirmed using polymerase chain reaction analysis of DNA extracts from ears [31]. After one week of acclimatization, they were fed on HCHF diet containing 15% lard and 0.5% cholesterol (Biotech company, China) for eight weeks. The Ldlr-/- hamsters were then randomly divided into three groups according to their weights (n=8 per group) and orally administered with a mixture of volume vehicle (distilled water), tetramethylpyrazine (32mg/kg/d) and clopidogrel (32mg/kg/d) drugs for eight weeks. Wild type golden Syrian hamsters (n=8) purchased from Vital River Laboratory (Charles River, Beijing, China) were fed on a standard chow diet as healthy control. All hamsters were maintained on a 12-hour light/12-hour dark cycle with free access to water.

Finally, the hamsters were fasted for 12h and anesthetized through intraperitoneal injection of 1% sodium phenobarbital (70mg/kg). Blood samples were taken from abdominal aortas, and plasma was separated by centrifugation for 10 min at 2700×g. TC, TG, and HDL were determined using commercially available kits (BIOSINO, China).

#### Oil red O staining

As described previously [31][44], anesthetized hamsters were perfused with 0.01M PBS through the left ventricle. In brief, hearts and whole aortas were placed in 4% paraformaldehyde solution overnight and transferred to 20% sucrose solution for one week. Hearts were then fixed into OCT compound and cross-sectioned (8 um per slice). The atherosclerotic lesions in aortic root were stained with 0.3% Oil red O solution (Solarbio, China), rinsed with 60% isopropanol and distilled water and counterstained with hematoxylin. The results were represented by the percentage positive area of total area (en face analysis) and net lesion area (aortic root sections). Images were analyzed with Image J [45].

#### Immunohistochemistry analysis

Analysis of atherosclerotic plaque cell composition was determined by immunohistochemistry analysis of the aortic root. Macrophages and SMC were stained with CD68 (BOSTER, BA36381:100) antibody and a-SMA antibody (BOSTER, A03744, 1:100), as reported previously in hamster researches [31]. Then biotinylated second antibody (Vector Laboratories, ABC Vectastain, 1:200) were used for incubation under 2% normal blocking serum. The cryosections were visualized using 3,3-diaminobenzidine (Vector Laboratories, DAB Vectastain). The results were represented by the percentage positive area of the total cross-sectional vessel wall area in the aortic root sections and analyzed using Image J [45].

#### Washed platelet preparation

Blood per hamster, 3 to 4 mL was collected from abdominal aortas into a tube containing an acid-citrate-dextrose anticoagulant (83.2mM D-glucose, 85mM trisodium citrate dihydrate, 19mM citric acid monohydrate, pH5.5). Platelet-rich plasma (PRP) was prepared after centrifugation at 300×g for 10min in room temperature. For washed platelet preparation, PRP was centrifuged at 1500×g for 2min. After collecting supernatant consisting of platelet-poor plasma into another centrifuge tube, the remaining PRP was washing three times, and the pellet was re-suspended in a modified Tyrode buffer (2.4mM HEPES, 6.1mM D-glucose, 137mM NaCl, 12mM HaHCO3, 2.6mM KCl, pH7.4).

#### Assessment of platelet activity

Washed platelets were loaded with fura-2/AM(5μM, Molecular Probe) in the presence of Pluronic F-127 (0.2μg/mL, Molecular Probe) and then incubated at 37□ for 1 hour in the dark [46]. Platelets were washed and re-suspended in Tyrode buffer containing 1mM calcium. After activation of ADP (20μM, Sigma), intracellular calcium concentration was measured using a fluorescence mode of Synergy H1 microplate reader (Biotek, USA). Excitation wavelengths was alternated at 340 and 380 nm. Excitation was measured at 510 nm. TritonX-100 and EGTA were used for calibration of maximal and minimal calcium concentrations, respectively. Washed platelets were activated by ADP and then lysed by 0.1M HCl on ice. According to the manufacturer’s instructions, the level of intracellular cAMP was determined by ELISA (Enzo Life Sciences, ADI-900-066).

#### Western blot analysis

Washed platelets from each group were lysed with radioimmunoprecipitation assay (RIPA) buffer with the presence of protease and phosphatase inhibitor mixtures on ice (Solarbio, China). Lysates were separated by 10000×g centrifugation for 10 min at 4□. Total protein concentrations were determined by BCA method. Equal amounts of total protein (40μg) were resolved in SDS-PAGE and electroblotted. The nitrocellulose membranes were blocked with 5% skimmed milk at room temperature for 2 hours and incubated with primary antibodies targeting PI3K(CST, 4257T, 1:500), Akt(CST, 9272, 1:2000), p-Akt(CST,2965,1:1000) and GADPH (Abcam, ab8245, 1:5000) overnight at 4□. The membranes were then incubated with the HRP-conjugated anti-rabbit antibody for 1 hour at 37□, followed by enhanced chemiluminescence detection.

### Statistical analysis

All data were expressed as mean ±standard error (SEM). Shapiro-Wild test and Levene’s test were used for determining normality of data distribution and homogeneity of variances, respectively. An unpaired student’s t-test was used to compare data among different groups when data were normally distributed, and variances were equal among the groups. Unpaired t test with Welch’s correction was used when there was unequal standard deviation among groups. Mann-Whitney test was used for nonparametric test. All p-values less than 0.05 were considered statistically significant. All statistical analyses were performed using GraphPad Prism 8.0 (GraphPad, United states).

## Author contributions

Y.Z. and D.Z.S. spearheaded and supervised all the experiments. Y.Z., D.Z.S., Y.H.Z. and K.Z. designed research. Y.H.Z, K.Z., B.Y.G., L.S., M.M.G., Y.H.W., and J.G. conducted experiments. L.S. and B.Y.G. analyzed data. Y.H.Z, K.Z., and Y.Z. prepared the manuscript. All authors reviewed and approved the manuscript.

## Disclosure of Potential Conflicts

The authors declare that none of them have any conflict of interest.

## Acknowledgements

This work was funded by the National Natural Science Foundation of China, grant (No. 81703927) and the Fundamental Research Funds for the Central public welfare research institutes of China, grant (No. ZZ13-YQ-008). The images of atherosclerosis and platelets in the graphical abstract were adapted from Servier Medical Art (http://smart.servier.com). Dr. Jerry, a professional English editor, provided language help and writing assistance.

## Notes

https://github.com/CUMTzackGit/DLDTI

### Competing Interest Statement

The authors have declared no competing interest.

